# Restoration of Rapid-Eye Movement Sleep During Cocaine Abstinence Reduces Incubation of Cocaine Seeking and Normalizes Dopamine Transporter Function

**DOI:** 10.64898/2026.01.12.697196

**Authors:** IP Alonso, SR Cohen, VM Migovich, RA España

## Abstract

Progressive increases in cocaine craving or seeking during abstinence from cocaine are thought to contribute to relapse. Recent observations suggest that sleep disruptions during abstinence from cocaine may have a significant influence on cocaine seeking. While the neural mechanisms underlying the association between sleep disruptions and increases in cocaine seeking during abstinence continue to be investigated, alterations in mesolimbic dopamine transmission may be a common factor. In these studies, we assessed whether sleep disruptions during abstinence are associated with incubation of cocaine seeking and dopamine terminal adaptations in the nucleus accumbens core of female and male rats. We observed that intermittent access to cocaine followed by abstinence reduced rapid-eye movement sleep, intensified cue-induced cocaine seeking, and enhanced dopamine uptake and dopamine transporter sensitivity to cocaine. Notably, a sleep restoration procedure that restricted sleep to the light phase (active period) restored rapid-eye movement sleep, prevented incubation of cocaine seeking, and normalized baseline dopamine uptake and dopamine transporter sensitivity to cocaine. These findings indicate that rapid-eye movement sleep disruptions during abstinence contribute to exaggerated cocaine seeking and that dopamine transporter adaptations are a potential molecular substrate through which these changes occur. Thus, interventions that restore sleep during cocaine abstinence may serve as effective behavioral therapies for reducing cocaine craving and preventing subsequent relapse.

## INTRODUCTION

A progressive intensification, or ‘incubation’, of cocaine craving has been demonstrated to occur during abstinence from drugs of abuse including cocaine (Parvaz et al, 2016). The cocaine craving phenomenon has been modeled repeatedly in rodents using cue-induced cocaine seeking tests and has been hypothesized to contribute to relapse to cocaine use (Garavan et al., 2000; Grimm et al., 2001; Puhl et al., 2013; Angarita et al., 2014; Chen et al., 2015; Alonso et al., 2022). Although the factors that influence incubation of cocaine craving and seeking remain incompletely understood, recent observations suggest that sleep disruptions during abstinence may contribute to incubation of cocaine seeking (Chen et al., 2015; Angarita et al., 2016; Bjorness and Greene, 2021). Indeed, both clinical and animal studies indicate significant sleep disruptions during abstinence from cocaine, which often manifest as reduced rapid-eye movement (REM) sleep and fragmented sleep architecture (Dugovic et al., 1992; Morgan and Malison, 2007; Matuskey et al., 2011; Yang et al., 2011; Roncero et al., 2012; Angarita et al., 2014; Chen et al., 2015; Garcia and Salloum, 2015; Parvaz et al., 2016; Bjorness and Greene, 2021). While the neural mechanisms underlying the association between REM sleep disruptions and incubation of cocaine seeking continue to be studied, accumulating evidence suggests that alterations in mesolimbic dopamine transmission may be a common factor in each of these processes (Woolverton and Johnson, 1992; Wise et al., 1996; Koob and Le Moal, 1997; Phillips et al., 2003; Calipari et al., 2015; Siciliano et al., 2015; Brodnik et al., 2020b).

Recent findings suggest that dopamine transporter (DAT) alterations during abstinence from cocaine may contribute to progressive increases in cocaine seeking and motivation. For example, we demonstrated that intermittent access (IntA) to cocaine promoted: 1) robust cocaine seeking and motivation both early and later in abstinence; 2) enhanced dopamine uptake rate in the nucleus accumbens (NAc) core following abstinence; and 3) increased DAT sensitivity to cocaine (i.e. inhibition of dopamine uptake) in the NAc core (Alonso et al., 2022; Clark et al., 2024; Samels et al., 2024). Others report similar findings, demonstrating that IntA to cocaine promotes motivation for cocaine, enhances dopamine uptake rate, and increases DAT sensitivity after abstinence from IntA to cocaine (Calipari et al., 2013; Calipari et al., 2015). Together, these findings suggest that IntA to cocaine promotes aberrant DAT adaptations that persist throughout abstinence and coincide with increased cocaine seeking.

Further evidence suggests that mesolimbic dopamine participates in the regulation of sleep/wake behavior. For example, ventral tegmental area (VTA) dopamine neurons displayed higher activity during REM sleep compared to wakefulness (WAKE) and non-REM (NREM) sleep (Dahan et al., 2007; Eban-Rothschild et al., 2016). Activation of VTA dopamine neurons initiates and maintains wakefulness even after hours of sleep deprivation, and this effect is primarily mediated by projections to the NAc (Eban-Rothschild et al., 2016; Oishi et al., 2017). Our prior work also demonstrated that the DAT governs diurnal fluctuations in extracellular dopamine tone in the NAc (Ferris et al., 2014), and that dopamine uptake rate and dopamine transporter phosphorylation at Thr53 (pDAT) vary across sleep/wake behavior (Alonso et al., 2021). Given that variations in dopamine uptake and pDAT expression influence DAT sensitivity to cocaine, cocaine intake, and motivation for cocaine (Calipari et al., 2017; Challasivakanaka et al., 2017; Brodnik et al., 2020b; Alonso et al., 2021; Shaw et al., 2021; Clark et al., 2024). These observations support the hypothesis that sleep disruptions during cocaine abstinence contribute to DAT adaptations that ultimately influence cocaine seeking.

Despite these observations, it remains unknown whether IntA to cocaine—which mimics the clinically reported binge pattern of cocaine consumption—generates sleep disruptions, and the extent to which these processes involve adaptations in DAT function. In the current studies, we used a combination of sleep/wake recordings, IntA to cocaine self-administration, behavioral sleep manipulations, and cue-induced seeking tests to examine to what degree cocaine seeking is associated with sleep disruptions. We further assessed whether these effects involve changes in dopamine transmission using fast scan cyclic voltammetry (FSCV). We hypothesized that IntA to cocaine disrupts sleep/wake architecture during abstinence and that normalizing this aberrant sleep/wake activity would subsequently reduce both cue-induced cocaine seeking and aberrant dopamine transmission. Rats underwent baseline electroencephalographic (EEG) and electromyographic (EMG) sleep/wake assessments, followed by 7 days of self-administration with IntA to cocaine and 7 days of abstinence. Cocaine craving was assessed with cue-induced seeking tests on abstinence days 1 and 7, with continuous sleep/wake recordings throughout this period. During abstinence, rats were either undisturbed or subjected to a sleep restoration procedure adapted from a prior report (Chen et al., 2015). The day following the last seeking test, brain slices containing the NAc were prepared to assess dopamine transmission using FSCV.

## METHODS

### Subjects

Adult female (200-225g) and male Long-Evans rats (325-350g; Envigo, Frederick, MD, USA) were maintained on a 12-hour reverse light/dark cycle (lights on at 15:00; lights off at 03:00) and given ad libitum access to food and water. After arrival, rats were given 7 days to acclimate before surgery. All protocols and animal procedures were conducted in accordance with the National Institutes of Health Guide for the Care and Use of Laboratory Animals under the supervision of the Institutional Animal Care and Use Committee at Drexel University College of Medicine.

### Surgical Procedures

Rats were anesthetized using 2.5% isoflurane and were implanted with a silastic catheter (ID, 0.012 in OD, 0.025 in. Access Technologies, Skokie, IL) placed in the right jugular vein for intravenous delivery of cocaine. The catheter was connected to a cannula port which exited through the skin on the dorsal surface in the region of the scapulae. Within the same surgery, rats were next placed into a stereotaxic frame and implanted with electroencephalographic (EEG) and electromyographic (EMG) electrodes for sleep/wake recordings. For recording EEG activity, one screw electrode (Plastics One, Roanoke, VA, USA) was implanted above the frontal cortex (+1.3 mm A/P, + 1.3 mm M/L, and -1.0 mm D/V), and a second electrode was implanted ipsilaterally over the hippocampus (+2.4 mm A/P, +3.2 mm M/L, and -1.0 mm D/V). A third screw electrode was implanted in the contralateral cerebellum (+1.3 mm A/P, - 1.3 mm M/L, and -1.0 mm D/V) to serve as a ground. Two EMG wire electrodes were threaded into the dorsal neck muscle for recording muscle activity. EMG electrodes consisted of 100 mm insulated stainless-steel wires (Cooner Wire, Charsworth, CA, USA) with 2 mm of exposed wire in contact with muscle. All electrodes were routed through a 6-pin connector (Plastics One) and cemented (Jet Liquid with acrylic denture repair powder) into place on the skull. Ketoprofen (Patterson Veterinary, Devens, MA; 5mg/kg s.c. of 5 mg/ml) and Enrofloxacin (Norbrook, Northern Ireland; 5 mg/kg s.c. of 5 mg/ml) were provided at the time of the surgery and again 12 h later. In addition, antibiotic/analgesic powder (Neopredef, Kalamazoo, MI) and skin glue (VetBond™, 3M, St. Paul, MN) were applied around the head, chest, and back incisions. Rats were subsequently singly housed and allowed to recover for 7 days before sleep recordings began. Intravenous catheters were manually flushed daily with 0.5 ml of gentamicin antibiotic (Aspen Veterinary Resources, Greeley, CO; 5 mg/kg i.v. of 5 mg/ml) dissolved in heparinized saline (Meitheal Pharmaceuticals, Chicago, IL; 20 U/ml) to prevent infection and catheter occlusion.

### Sleep Recordings

Rats were placed in a custom-designed testing chamber, which consisted of an open-top, plexiglass cylinder (30 cm in diameter, 38 cm high) with a rotating bar attached parallel to the floor. Sleep chambers were housed in a sound-attenuated outer chamber containing an LED light (lights on at 15:00; lights off at 03:00) and a fan. Rats were supplied with food and water available ad libitum and connected to recording lines via a commutator which allowed unrestricted movement. EEG (0.3 – 100 Hz bandpass) and EMG signals (1 – 50 Hz bandpass) were amplified, filtered, and recorded using PowerLab hardware and LabChart software (AD Instruments, Inc., Colorado Springs, CO, USA) and analyzed using Sirenia SleepPro (Pinnacle Technology, Inc., Lawrence, KS, USA). Data were scored manually for analysis of the number of bouts, bout length, and percentage of time spent in wakefulness (WAKE), non-REM (NREM) sleep, and REM sleep, as previously described (Berridge and España, 2000; España et al., 2001; Brodnik et al., 2015; Alonso et al., 2021). Briefly, EEG signals were scored in 10 s bins (epochs) with NREM sleep defined as high-voltage EEG consisting of predominant delta frequency and low-voltage EMG, REM sleep defined as low-voltage EEG consisting of predominant theta frequency and minimal to no EMG activity, and WAKE defined as low-voltage EEG consisting of minimal delta and theta frequencies with mixed higher range frequencies and EMG activity of an average amplitude twice that observed in NREM. To be scored as a distinct bout, EEG and EMG activity patterns were required to persist for three consecutive 10 s epochs. After surgical recovery, baseline sleep/wake activity was recorded prior to cocaine exposure to serve as a baseline for within-subjects analysis. Rats first acclimated to the sleep recording chamber for a 24-h period before EEG/EMG signals were recorded for the subsequent 24 h to monitor WAKE, NREM and REM sleep before cocaine exposure. In addition to these baseline recordings, sleep/wake activity was also monitored throughout abstinence between seeking tests (see below).

### Self-Administration

After baseline sleep/wake activity recordings, rats self-administered cocaine in a (23.5 ✕ 15 ✕ 14 inches) operant chamber (Med Associates, St Albans, VT) located inside a sound-attenuating cabinet. A ventilating fan masked background noise. Two levers were located 6 cm above the grid floor on the left and right side of the left wall, the right lever was designated as active and the left one as inactive. A house light was located at the top center of the left wall. Pressing the active lever resulted in an intravenous infusion of cocaine hydrochloride (National Institute on Drug Abuse Drug Supply program) dissolved in 0.9% sterile saline. All measures were recorded using Med Associates software and began between 08:30-10:30, during the second half of the 12-h dark phase (03:00-15:00). Rats were first trained to self-administer cocaine on a 6-h fixed ratio 1, long access (LgA) schedule during which single active lever presses initiated an intravenous injection of cocaine (0.5 mg/kg, infused over 2.5 - 4 s) paired with a cue light. Acquisition was defined as a rat reaching ≥40 infusions in two consecutive sessions. Each rat underwent one cocaine self-administration session per day, for an average of 5 days per week. However, rats always self-administered cocaine the day before cue-induced seeking tests.

*Intermittent Access*: Once self-administration behavior was acquired, rats self-administered cocaine on the intermittent access (IntA) schedule (Zimmer et al., 2012; Calipari et al., 2013; Alonso et al., 2022). During the 6-h IntA sessions, rats had access to cocaine for 5-min trials followed by 25-min timeout periods when the levers were retracted. Active lever presses resulted in one infusion of 0.375 mg/kg cocaine infusion over 0.9 s, paired with a cue light. At the start of each infusion, the stimulus light above the active lever was illuminated for the length of the infusion. There was no timeout period after infusions to allow a binge-like pattern of consumption. However, active lever presses that occurred during the 0.9 s cocaine delivery were recorded despite having no consequence.

### Cue-Induced Seeking Tests

To assess incubation of cocaine seeking, rats performed cue-induced cocaine seeking tests on abstinence day 1 (AD1) and abstinence day 7 (AD7). During the 1-h seeking tests, a single active lever press resulted in presentation of the cue and house lights that had been previously paired with cocaine, but no cocaine was delivered. Under these conditions, the number of active lever presses served as a measure of cocaine seeking, and the number of inactive lever presses served as a measure of nonspecific behavior.

### Abstinence

Following the AD1 seeking test, rats were randomly assigned to the IntA only control group (IntA) or the IntA with sleep restoration (IntA+SR) group (see below). During this phase, sleep/wake activity was recorded to assess sleep disruptions after abstinence. Immediately following the AD1 seeking test, rats were transferred to sound attenuated sleep chambers and connected to EEG/EMG recording cables. Rats were housed in the sleep chamber for the duration of abstinence until the AD7 seeking test and were returned to their home cages after the seeking test was completed.

### Sleep Restoration

Rats are nocturnal animals, with approximately 60% of their total daily sleep occurring during the light phase and approximately 60% of their activity occurring during the dark phase (Borbely, 1977; Neuhaus and Borbely, 1978; Alonso et al., 2021). Sleep homeostatic regulation consists of increased sleep need following extended wakefulness (Neuhaus and Borbely, 1978; Borbely and Achermann, 1999; Hasler et al., 2012). Consistent with these findings, animal studies have demonstrated that inhibiting scattered “napping” (i.e., brief periods of sleep) during the dark phase, which is the normal active phase in rodents, improves REM sleep during the light/inactive phase, effectively constituting a form of ‘sleep restoration’ (Puhl et al., 2013; Chen et al., 2015). Sleep restoration for the IntA+SR rat group was implemented with a slowly rotating bar (5 rpm) at the bottom of the sleep chamber during the 12-h dark phase to encourage wakefulness. The bar moved constantly during the 12 h of the dark phase, changing rotation direction every 30 min, and remained stationary during the 12-hour light phase. By imposing wakefulness during the dark-phase, this sleep restoration approach results in accumulation of sleep pressure, thereby enhancing sleep quality by consolidating sleep to the inactive light period only (Chen et al., 2015). The bar rotated daily during the 12-h dark phase throughout the abstinence period except for the dark phase immediately prior to the seeking test on AD7 to assess the cumulative effects on AD7 sleep/wake activity and to avoid any interference between the acute effects of sleep restoration prior to the seeking test. The IntA group was housed identically to the IntA+SR group except that the rotation bar remained stationary the entirety of abstinence, thereby not manipulating sleep.

### *Ex vivo* fast scan cyclic voltammetry

Eighteen hours after the AD7 seeking test, rats were anesthetized with 2.5% isoflurane for 5 min, brains were rapidly dissected and transferred to ice-cold, oxygenated artificial cerebrospinal fluid (aCSF) containing NaCl (126 mM), KCl (2.5 mM), NaH2PO4 (1.2 mM), CaCl2 (2.4 mM), MgCl2 (1.2 mM), NaHCO3 (25 mM), glucose (11 mM), and L-ascorbic acid (0.4 mM), with pH adjusted to 7.4. A vibrating microtome was used to produce 400µm-thick coronal sections containing the NAc core. Slices were transferred to room-temperature oxygenated aCSF and left to equilibrate for 1 h before being transferred into a recording chamber flushed with aCSF (32°C, ∼150-200mL/min).

A bipolar stimulating electrode was placed on the surface of the tissue in the NAc core, and a carbon fiber electrode was placed between the stimulating electrode leads. Dopamine release was evoked every 3 min using a single electrical pulse (400µA; 4ms; monophasic), measured, and analyzed using Demon Voltammetry and Analysis Software (Yorgason et al., 2011). Once baseline dopamine release was stable (3 successive stimulations within <10% variation), the slice was exposed to gradual increases in cocaine concentrations (0.3 - 30µM) as previously described (Brodnik et al., 2020a; Alonso et al., 2021; Cohen et al., 2025). Dopamine concentrations were calculated by comparing currents at the peak oxidation potential for dopamine in consecutive voltammograms with electrode calibrations determined using an *in situ* calibration method as described previously (Roberts et al., 2013; Meunier et al., 2017; Brodnik et al., 2020a; Alonso et al., 2021). To determine influences on dopamine transmission, we assessed the amplitude of stimulated dopamine release (dopamine peak height), maximal dopamine uptake rate (*Vmax*; dopamine uptake), and cocaine-induced inhibition of dopamine uptake (app *Km*) using a Michaelis-Menten based model (Prince et al., 2015; Levy et al., 2017; Brodnik et al., 2020a). Baseline dopamine uptake was determined by setting Km values to 0.18 µM and all cocaine-induced alterations in uptake were attributed to changes in apparent *Km*. Inhibition constants (*Ki*) were determined to calculate the necessary cocaine concentration to produce 50% of app *Km* and were then calculated using the equation [*Km* / slope] (Alonso et al., 2021). Controls for FSCV experiments were cocaine-naive rats housed in the same room as rats undergoing self-administration.

### Statistical Analyses

Data were analyzed using Prism GraphPad 10. Each rat served as its own within-subject control for comparing sleep across days of abstinence and active lever presses across seeking tests. Self-administration data prior to sleep restoration were analyzed using Student’s t-tests or mixed design ANOVAs with group as the between-subjects variable (future IntA versus IntA+SR) and IntA session (1-7) as the within-subjects variable. Sleep/wake measures (percentage of time, bout number, and average bout duration) on AD1 and were analyzed using mixed design ANOVAs with group as the between-subjects variable (IntA vs IntA+SR) and day (baseline versus AD1) as the within-subjects variable for the 12-h dark phase. Cue-induced cocaine seeking was analyzed using a mixed design ANOVA with group as the between-subjects variable (IntA versus IntA+SR) and day (AD1 versus AD7) as the within-subjects variable. Sleep/wake measures (percentage of time, bout number, and average bout duration) on AD7 were analyzed using mixed design ANOVAs with group as the between-subjects variable (IntA vs IntA+SR) and day (baseline versus AD7) as the within-subjects variable for the 12-h light phase. Baseline FSCV data for dopamine peak height and dopamine uptake rate were analyzed using one-way ANOVAs with group as the between-subjects variable (naive, IntA, IntA+SR). FSCV measures of dopamine peak height and DAT sensitivity to cocaine following increasing concentrations of cocaine were analyzed using mixed two-way ANOVAs with group as the between-subjects variable (IntA vs IntA+SR) and cocaine concentration as the within-subjects variable (0.3, 1, 3, 10, and 30 µM). *Ki* was analyzed using a one-way ANOVA with group as the between-subjects variable (naive, IntA, IntA+SR). Šídák’s multiple comparison post-hoc tests were used following significant mixed design ANOVAs in the self-administration and sleep experiments while Dunnett’s multiple comparison post-hoc tests were used following significant one-way ANOVAs or mixed design ANOVAs for the FSCV experiments. Pearson correlations were used to assess if the percentage of WAKE, NREM, or REM on AD7 relative to baseline sleep/wake recordings predicted lever pressing on AD7, dopamine uptake rate, or *Ki*.

## RESULTS

### Cocaine intake and lever pressing were comparable prior to the IntA and IntA sleep-restored group assignment

To assess the effects of cocaine and sleep restoration on sleep, incubation of cue-induced seeking, and DAT function in the NAc during abstinence, rats first self-administered cocaine on an IntA schedule (**Fig. 1A**). We observed that there were no significant differences in self-administration behavior between rats that would later be separated into the IntA and IntA+SR groups. Student’s t-tests demonstrated no differences between groups for days to acquire (*t*_(19)_ = 1.420, *p*=0.1719; **Fig. 1B**) or total cocaine intake across the acquisition (LgA) and IntA portions of self-administration (*t*_(19)_ = 0.2741, *p*=0.7870; **Fig. 1C**). A mixed design ANOVA with group as the between-subjects variable (IntA vs IntA+SR) and self-administration session (1-7) as the within subjects variable revealed no effect of group (*F*_(1,19)_ = 0.0899, *p*=0.7675) or session (*F*_(4.403, 83.66)_ = 0.2632, *p*=0.9150), and no group × session interaction on cocaine intake during the IntA portion of self-administration (*F*_(4.403, 83.66)_ = 0.5325, *p*=0.7290; **Fig. 1D**). Further, a mixed design ANOVA for infusions across IntA session revealed no effect of group (*F*_(1,19)_ = 0.5417, *p*=0.4707) or session (*F*_(3.897,74.05)_ = 0.2458, *p*=0.9075), and no group × session interaction (*F*_(3.897,74.05)_ = 0.6361, *p*=0.6344; **Fig. 1E**). Similarly, a mixed design ANOVA for active lever presses showed no effect of group (*F*_(1,19)_ = 0.5376, *p*=0.4724), session (*F*_(3.816,72.50)_ = 0.1098, *p* = 0.9755), or a group × session interaction (*F*_(3.816,72.50)_ = 0.7709, *p*=0.5421; **Fig. 1F**). Lastly, a mixed design ANOVA for inactive lever presses revealed no effect of group (*F*_(1,19)_ = 0.3590, *p*=0.5559), session (*F*_(1.086,20.64)_ = 0.5071, *p*=0.4992), or a group × session interaction (*F*_(1.086,20.64)_ = 1.004, *p*=0.3353; **Fig1. G**).

**Figure 1.**
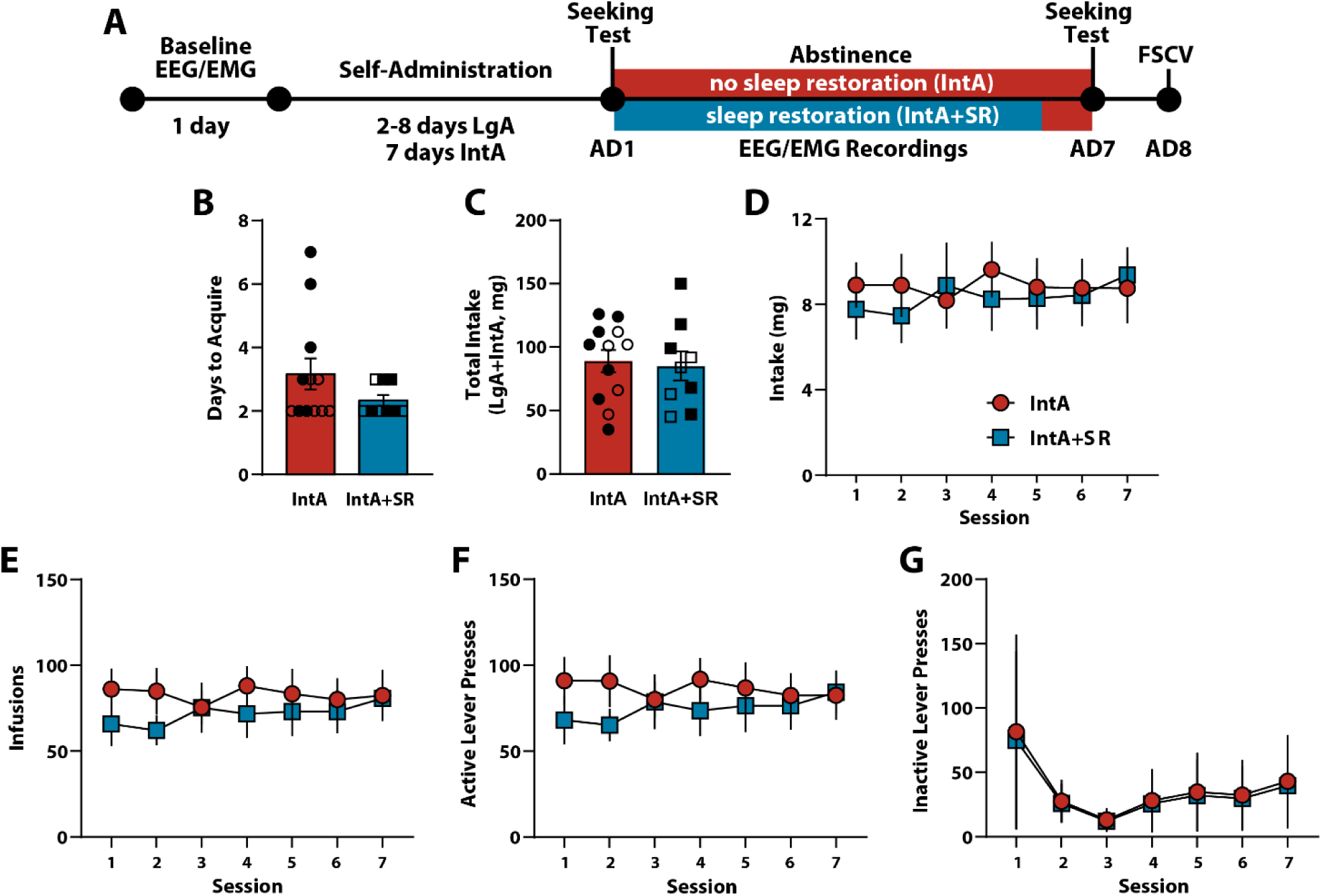
Experimental timeline and self-administration. **(A)** Experimental timeline. **(B)** Number of days to acquire LgA self-administration and **(C)** total cocaine intake during IntA self-administration. **(D)** Cocaine intake, **(E)** cocaine infusions, **(F)** active lever presses, and **(G)** inactive lever presses across IntA sessions. Data are presented as mean ± S.E.M. ○□females, ●◼ males. IntA n=12, IntA+SR n=9.

### Sleep restoration increased wakefulness and reduced sleep during the active/dark phase

To validate that the sleep restoration procedure limits sleep during the dark phase, we examined the first exposure to the rotating bar for the IntA+SR group during the 12-hr dark period on AD1 (during the 24 hrs after the AD1 seeking test). We then analyzed the percentage of time spent in a given sleep/wake state (i.e. WAKE, NREM, REM), as well as total bout number and average bout duration for each sleep/wake state.

#### WAKE during the 12-hr dark period on AD1

IntA to cocaine reduced wakefulness, whereas sleep restoration using the continuously rotating bar during the dark period significantly increased time spent awake. A mixed design ANOVA with group (IntA vs IntA+SR) as the between-subjects variable and day (baseline vs AD1) as the within-subjects variable revealed a significant effect of group (*F*_(1,17)_ = 71.49, *p*<0.0001) and a significant day × group interaction (*F*_(1,17)_ = 26.38, *p*<0.0001), but no effect of day (*F*_(1,17)_ = 0.03265, *p=*0.8587) on the percentage of time spent in WAKE (**Fig. 2A**). Šídák’s post-hoc tests demonstrated that compared to baseline activity, IntA to cocaine (*p*=0.0008) significantly decreased WAKE on AD1, while sleep restoration significantly increased WAKE (*p*=0.0125) on AD1 compared to baseline. We did not observe differences in WAKE bout number and WAKE bout duration. A mixed design ANOVA indicated no effect of group (*F*_(1,17)_ = 2.059, *p=*0.1695), day (*F*_(1,17)_ = 0.3533, *p=* 0.5601), and a trend for a day × group interaction (*F*_(1,17)_ = 3.236, *p=*0.0898) on bout number (**Fig. 2B**). Similarly, a mixed design ANOVA revealed no significant effect of group (*F*_(1,17)_ = 3.047, *p=*0.0989) or day (*F*_(1,17)_ = 1.757, *p*=0.2025), and a trend for a day × group interaction (*F*_(1,17)_ = 3.056, *p*=0.0985) on WAKE bout duration (**Fig. 2C**).

**Figure 2.**
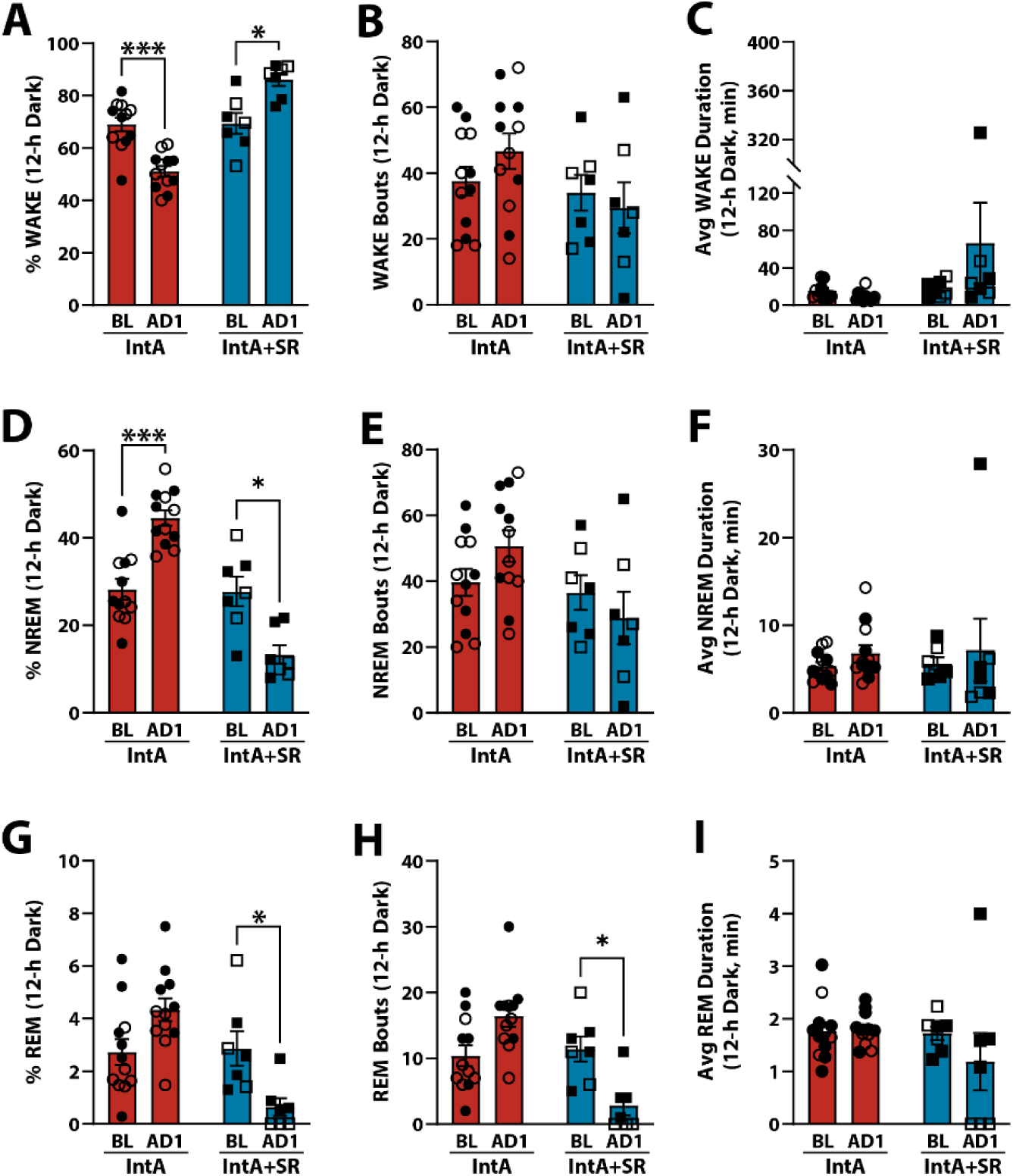
Sleep restoration reduced, dark-phase sleep early in abstinence. Percent of time spent in **(A)** WAKE, **(D)** NREM, and **(G)** REM during the first 12-hr dark period on AD1. Number of bouts for **(B)** WAKE, **(E)** NREM, and **(H)** REM during the first 12-hr dark period on AD1. Average bout duration for **(C)** WAKE, **(F)** NREM, and **(I)** REM during the first 12-hr dark period on AD1. Data shown as mean ± SEM. ○□females, ●◼ males. IntA n=12, IntA+SR n=7. Šídák’s tests \**p*<0.05, \*\*\**p*<0.001.

#### NREM during the 12-hr dark period on AD1

Changes in WAKE observed with IntA to cocaine and sleep restoration were associated with disruptions in NREM sleep. IntA to cocaine increased NREM sleep, whereas sleep restoration significantly decreased time spent in NREM. A mixed design ANOVA with group (IntA vs IntA+SR) as the between-subjects variable and day (baseline vs AD1) as the within-subjects variable revealed a significant effect of group (*F*_(1,17)_ = 77.75, *p*<0.0001) and a significant day × group interaction (*F*_(1,17)_ = 26.70, *p*<0.0001), but no effect of day (*F*_(1,17)_ = 0.0941, *p=*0.7628) on the percentage of time spent in NREM (**Fig. 2D**). Šídák’s tests showed that IntA to cocaine significantly increased the percentage NREM sleep (*p*=0.0006) on AD1 relative to baseline and that sleep restoration significantly decreased NREM (*p=*0.0142). These effects of sleep restoration on time spent in NREM, were associated with modest influences on NREM bout number and bout duration. A mixed design ANOVA revealed a significant day × group interaction (*F*_(1,17)_ = 5.248, *p=*0.0350), but no effect of group (*F*_(1,17)_ = 3.585, *p*=0.0755) or day (*F*_(1,17)_ = 0.1618, *p*=0.6925) on NREM bouts (**Fig. 2E**). However, Šídák’s tests showed that IntA to cocaine did not significantly affect the number of NREM bouts between baseline and AD1 (*p*=0.4391), and that sleep restoration produced only a trend for decreased bouts (*p*=0.0792). Further, a mixed design ANOVA for NREM bout duration revealed no significant effects of group (*F*_(1,17)_ = 0.0355, *p=*0.8527) or day (*F*_(1,17)_ = 1.552, *p*=0.2297) and no day × group interaction (*F*_(1,17)_ = 0.0003, *p=*0.9862; **Fig. 2F**).

#### REM during the 12-hr dark period on AD1

The most notable impact of sleep restoration on AD1 was on the percentage REM sleep and REM sleep bout number. A mixed design ANOVA with group (IntA vs IntA+SR) as the between-subjects variable and day (baseline vs AD1) as the within-subjects variable revealed a significant effect of group (*F*_(1,17)_ = 14.26, *p=*0.0015) and a significant day × group interaction (*F*_(1,17)_ = 11.86, *p=*0.0031), but no significant effect of day (*F*_(1,17)_ = 0.3018, *p=*0.5899) on the percentage of time spent in REM sleep (**Fig. 2G**). Šídák’s tests indicated that IntA to cocaine produced a strong trend for increased REM sleep on AD1 compared to baseline (*p*=0.0572) and that sleep restoration decreased REM sleep (*p=*0.0442). In terms of REM bout number, a mixed design ANOVA indicated a significant effect of group (*F*_(1,17)_ = 21.01, *p*=0.0003) and a significant day × group interaction (*F*_(1,17)_ = 12.49, *p*=0.0025), but no effect of day (*F*_(1,17)_ = 0.3889, *p*=0.5411; **Fig. 2H**). Šídák’s tests further showed that IntA to cocaine produced a strong trend for increased REM bout number (*p*=0.056), while sleep restoration significantly reduced REM bout number on AD1 relative to baseline (*p*=0.036). Finally, a mixed design ANOVA revealed no effect of group (*F*_(1,17)_ = 2.018, *p=*0.1735) or day (*F*_(1,17)_ = 1.024, *p*=0.3258), and no day × group interaction (*F*_(1,17)_ = 1.310, *p*=0.2682) on REM bout duration (**Fig. 2I**).

### Sleep restoration normalized REM sleep disruptions following abstinence from intermittent access to cocaine

To assess to what degree sleep restoration during the active/dark period normalizes sleep/wake activity on AD7 following IntA to cocaine, we analyzed the 12-hr inactive, light period on the last day of abstinence. We observed that IntA to cocaine had the most robust effects at reducing REM sleep and that sleep restoration restored REM sleep to baseline levels (**Fig. 3**).

**Figure 3.**
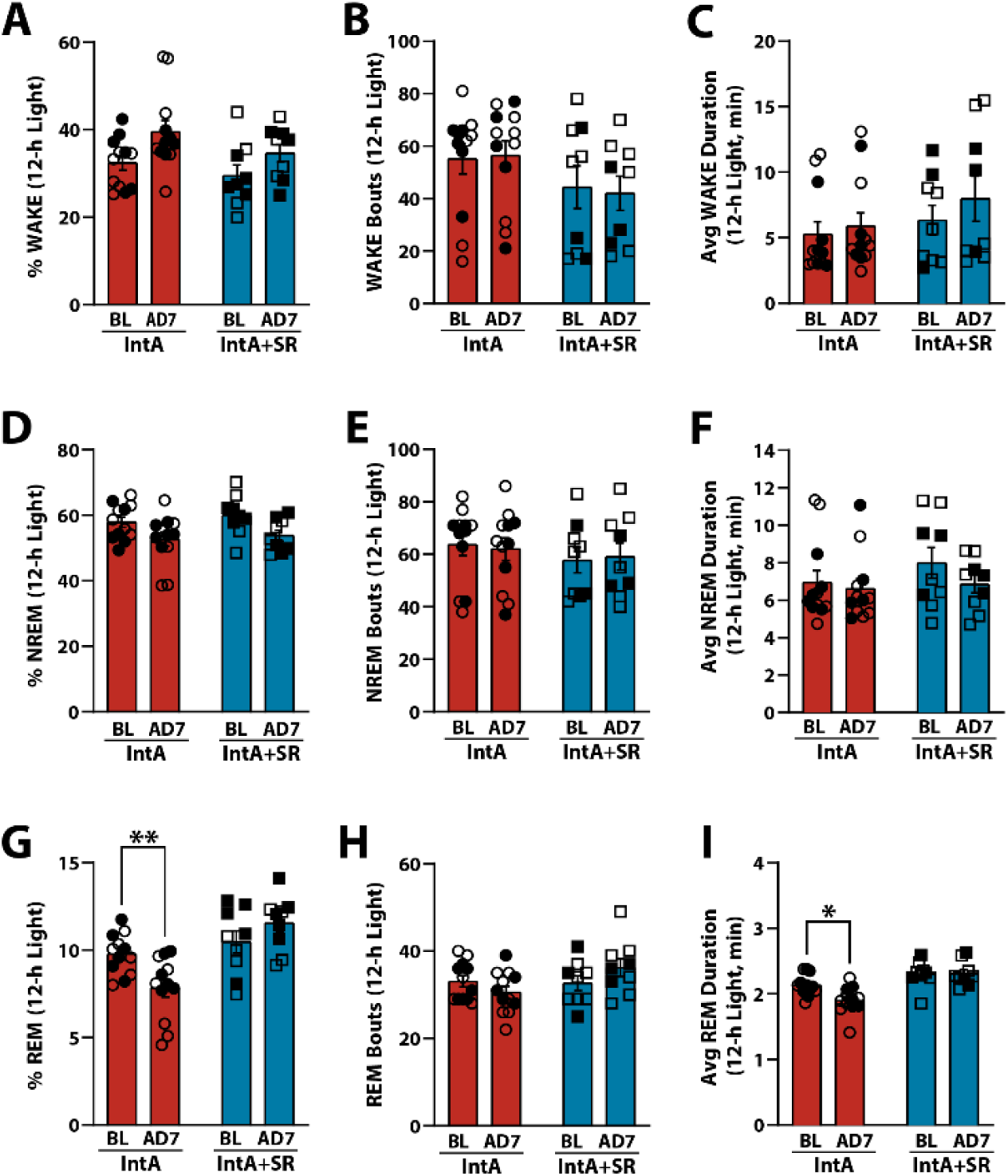
Sleep restoration normalized REM sleep disruptions observed following abstinence from intermittent access to cocaine. Percent of time spent in **(A)** WAKE, **(D)** NREM, and **(G)** REM during the 12-hr light period on AD7. Number of bouts for **(B)** WAKE, **(E)** NREM, and **(H)** REM during the during the 12-hr light period on AD7. Average bout duration for **(C)** WAKE, **(F)** NREM, and **(I)** REM during the 12-hr light period on AD7. Data shown as mean ± SEM. ○□females, ●◼ males. IntA n=12, IntA+SR n=9. Šídák’s tests \**p*<0.05, \*\**p*<0.01.

#### WAKE during the 12-hr light period on the last day of abstinence

Both IntA to cocaine alone and sleep restoration had only modest effects on WAKE activity on AD7 compared to baseline. A mixed design ANOVA with group (IntA vs IntA+SR) as the between-subjects variable and day (baseline vs AD7) as the within-subjects variable revealed a significant effect of day (*F*_(1,19)_ = 6.269, *p=*0.0216), but no significant effect of group (*F*_(1,19)_ = 3.677, *p=*0.0703) and no day × group interaction (*F*_(1,19)_ = 0.1657, *p*=0.6885) on the percentage of time spent in WAKE (**Fig. 3A**). Šídák’s tests showed that neither IntA alone (*p=*0.3302) nor sleep restoration (*p=*0.0756) had significant effects on the percentage of WAKE between AD7 and baseline, although there was a trend for increased WAKE with sleep restoration. IntA alone and sleep restoration also did not affect WAKE bout number or WAKE bout duration. A mixed design ANOVA showed no effect of group (*F*_(1,19)_ = 1.992, *p=*0.1743) or day (*F*_(1,19)_ = 0.0582, *p=*0.8119), and no group × day interaction (*F*_(1,19)_ = 0.6368, *p=*0.4347) on WAKE bout number (**Fig. 3B**). Moreover, a mixed design ANOVA of WAKE bout duration showed a significant effect of day (*F*_(1,19)_ = 5.018, *p=*0.0372), but no effect of group (*F*_(1,19)_ = 0.9579, *p*=0.3400) or a group × day interaction (*F*_(1,19)_ = 1.038, *p*=0.3211; **Fig. 3C**). Šídák’s tests showed that neither IntA alone (*p=*0.5938) nor sleep restoration (*p=*0.0864) had significant effects on WAKE bout duration on AD7 relative to baseline, though there was trend for increased WAKE bout duration following sleep restoration.

#### NREM during the 12-hr light period on the last day of abstinence

Similar to what was observed with WAKE, both IntA to cocaine alone and sleep restoration had only minor effects on NREM sleep on AD7. A mixed design ANOVA with group (IntA vs IntA+SR) as the between-subjects variable and day (baseline vs AD7) as the within-subjects variable showed a significant effect of day (*F*_(1,19)_ = 7.257, *p*=0.0144), but no effect of group (*F*_(1,19)_ = 0.9728, *p*=0.3364) or day × group interaction (*F*_(1,19)_ = 0.0579, *p*=0.8124) on the percentage of time spent in NREM sleep (**Fig. 3D**). Further, Šídák’s tests showed no significant effects of either IntA alone (*p=*0.1300) or sleep restoration (*p*=0.1471) on the percentage of NREM sleep between AD7 and baseline. There were also no effects of either IntA alone or sleep restoration on either NREM bout number or bout duration. A mixed design ANOVA revealed no effect of group (*F*_(1,19)_ = 0.5013, *p=*0.4875) or day (*F*_(1,19)_ = 0.0046, *p=*0.9465), and no group × day interaction (*F*_(1,19)_ = 0.5060, *p=*0.4855) on NREM bouts (**Fig. 3E**). Likewise, a mixed design ANOVA revealed no effect of group (*F*_(1,19)_ = 0.6280, *p=*0.4379) or day (*F*_(1,19)_ = 3.840, *p=*0.0649), and no group × day interaction (*F*_(1,19)_ = 1.082, *p=*0.3113) on NREM bout duration (**Fig. 2F**).

#### REM during the 12-hr light period on the last day of abstinence

IntA to cocaine significantly reduced REM sleep on AD7 while sleep restoration normalized these REM sleep disruptions. A mixed design ANOVA with group (IntA vs IntA+SR) as the between-subjects variable and day (baseline vs AD7) as the within-subjects variable demonstrated a significant effect of group (*F*_(1,19)_ = 14.53, *p*=0.0012) and a significant day × group interaction (*F*_(1,19)_ = 12.77, *p=*0.0020), but no effect of day (*F*_(1,19)_ = 1.050, *p*=0.3183) on the percentage of time spent in REM sleep (**Fig. 3G**). Šídák’s tests further indicated that IntA to cocaine significantly decreased the percentage of REM between AD7 and baseline (*p=*0.0047), and that sleep restoration prevented this decrease in REM (*p=*0.2048). These changes in time spent in REM sleep were related to changes in REM bout duration, but not REM bout number. A mixed design ANOVA for REM bouts revealed a significant group × day interaction (*F*_(1,19)_ = 5.920, *p=*0.0250), but no effect of group (*F*_(1,19)_ = 1.980, *p=*0.1755) or day (*F*_(1,19)_ = 0.1799, *p=* 0.6762; **Fig. 3H**). However, Šídák’s tests showed that neither IntA alone (*p=*0.2628) nor sleep restoration (*p=*0.1427) had significant effects on REM bout number. Finally, a mixed design ANOVA revealed a significant effect of group (*F*_(1,19)_ = 25.65, *p*<0.0001) but no effect of day (*F*_(1,19)_ = 3.401, *p=*0.0808) or group × day interaction (*F*_(1,19)_ = 3.797, *p=*0.0663) on REM bout duration (**Fig. 3I**). Šídák’s tests revealed that IntA alone (*p=*0.0184) significantly reduced REM bout duration on AD7 compared to baseline and that sleep restoration prevented this effect (*p=*0.9970).

### Sleep restoration prevented incubation of cue-induced cocaine seeking

Sleep disruptions during drug abstinence are a critical factor for promoting relapse, even after prolonged abstinence (Hasler et al., 2012; Dolsen and Harvey, 2017). Consistently, chronic sleep deprivation during cocaine self-administration increases the incentive value of cocaine and motivation for cocaine in rats (Puhl et al., 2013). To examine the impact of sleep restoration on incubation of cocaine seeking, we tested to what extent IntA to cocaine promoted active lever pressing during cue-induced seeking tests and if sleep restoration prevented this effect. Similar to our prior observations (Alonso et al., 2022; Clark et al., 2024), IntA to cocaine produced robust incubation of cocaine seeking as indicated by a significant increase in active lever pressing on AD7 compared to AD1. Sleep restoration prevented this incubation of cocaine seeking. A mixed design ANOVA with group as the between-subjects (IntA vs IntA+SR) and day (AD1 vs AD7) as the within-subjects variable revealed a significant effect of day (*F*_(1,19)_ = 28.84, *p*<0.0001), but no effect of group (*F*_(1,19)_ = 1.603, *p=*0.2207) and no group × day interaction (*F*_(1,19)_ = 3.559, *p=*0.0746; **Fig. 4A**). Šídák’s tests demonstrated that IntA significantly increased seeking from AD1 to AD7 (*p<*0.0001) while sleep restoration minimized this increase (*p*<0.0643). A mixed design ANOVA for inactive lever presses revealed no effect of group (*F*_(1,19)_ = 2.022, *p*=0.1712) or day (*F*_(1,19)_ = 0.8334, *p=*0.3727), and no group × day interaction (*F*_(1,19)_ = 1.578, *p=*0.2243; **Fig. 4B**). Together, these findings indicate that IntA to cocaine promotes robust incubation of cue-induced cocaine seeking and that sleep restoration prevents this increase in cocaine seeking.

**Figure 4.**
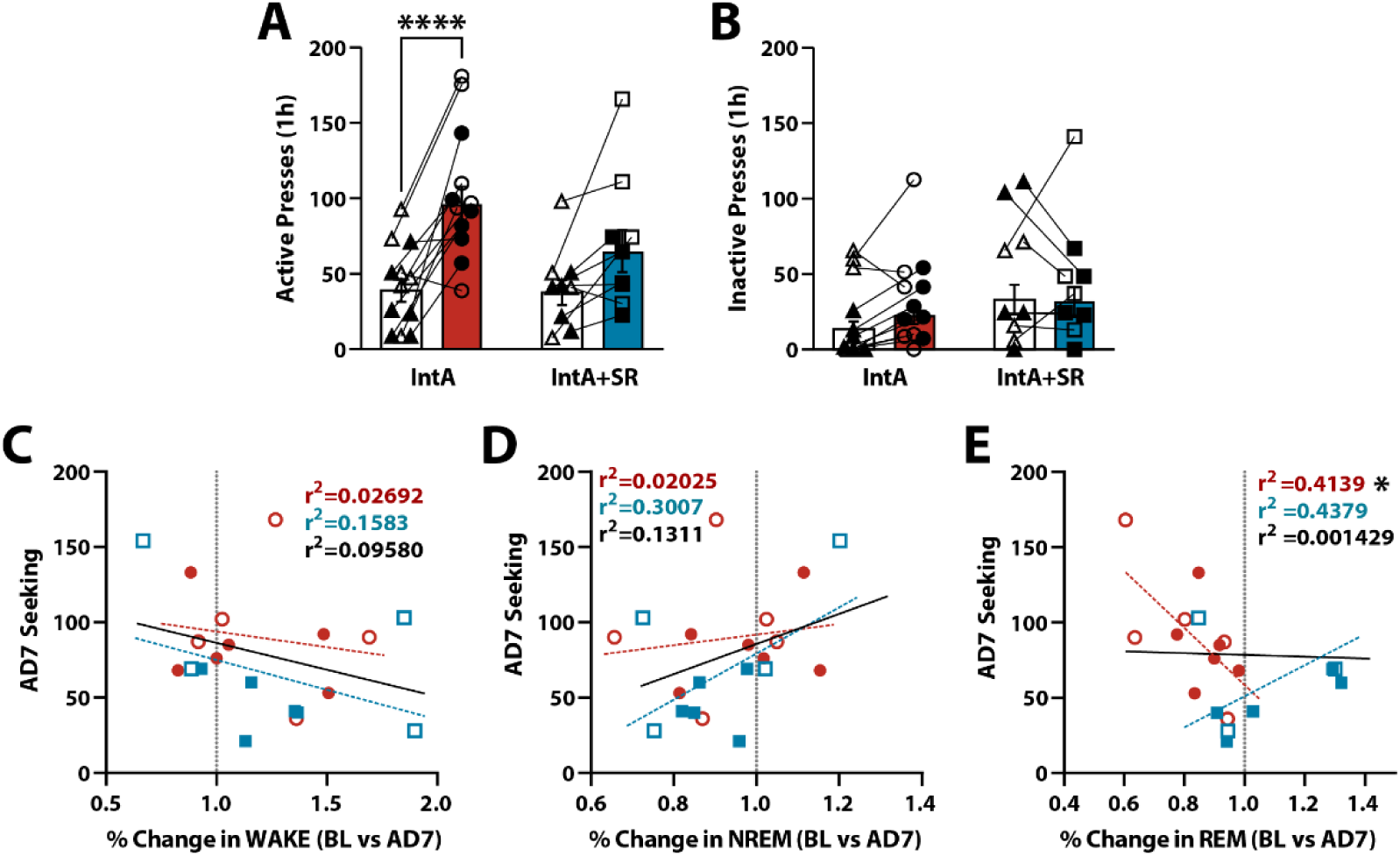
Sleep restoration reduced cue-induced cocaine seeking early in abstinence. **(A)** Active and **(B)** inactive lever presses during cue-induced seeking tests on AD1 and AD7. Pearson correlations between active presses on AD7 and the percent change from baseline (BL) in time spent in **(C)** WAKE, **(D)** NREM, or **(E)** REM. Black lines depict trendline for correlations with the IntA and IntA+SR groups combined. Vertical dashed lines indicate 100% of baseline. Data shown as mean ± SEM. ○□females, ●◼ males. IntA n=12; IntA+SR n=9. Šídák’s tests \*\*\*\**p*<0.00001. Pearson correlation \**p*<0.05.

We next examined if changes in sleep/wake behavior after IntA to cocaine predicted lever pressing on AD7. A Pearson correlation for all rats (IntA and IntA+SR) indicated no significant relationship between the percentage of change in WAKE (baseline vs AD7) and AD7 lever pressing (*F*_(1,18)_ = 1.907; R^2^ = 0.0958, *p*=0.1842; **Fig. 4C**). Likewise, we did not observe significant correlations between the percentage of change in WAKE and AD7 pressing for the IntA (*F*_(1,9)_ = 0.2490; R^2^ = 0.0269, *p*=0.6298), or the IntA+SR groups (*F*_(1,7)_ = 1.316; R^2^ = 0.1583, *p*=0.2889) when examined separately. We also observed no significant correlations b between the percentage of change in NREM and AD7 lever pressing when examining either all rats combined (*F*_(1,18)_ = 2.716; R^2^ = 0.1311, *p*=0.1167), the IntA group (*F*_(1,9)_ = 0.1860; R^2^ = 0.0203, *p*=0.6764), or the IntA+SR group (*F*_(1,7)_ = 3.010; R^2^ = 0.3007, *p*=0.1263) when examined separately (**Fig. 4D**). Lastly, when testing if baseline REM predicted AD7 lever pressing we observed no significant correlations when examining all rats combined (*F*_(1,18)_ = 0.0258; R^2^ = 0.0014, *p*=0.8743; **Fig. 4E**). However, there was a significant correlation between the percentage of change in REM and AD7 pressing for the IntA group (*F*_(1,9)_ = 6.355; R^2^ = 0.4139, *p*=0.0327) and a strong trend for significance for the IntA+SR group (*F*_(1,7)_ = 5.454; R^2^ = 0.4379, *p*=0.0522). Notably, all IntA alone rats exhibited a reduction in time spent in REM sleep during the AD7 12-h light phase (i.e. <100% of baseline; **Fig. 4E**). In contrast, sleep restoration increased time spent in REM sleep for several rats (i.e. >100% of baseline; **Fig. 4E**). These findings suggest that IntA not only promoted robust reductions in REM sleep after 7 days of abstinence but also promoted incubation of cue-induced cocaine seeking, while IntA+SR effectively normalized the effects of IntA alone.

### Sleep restoration during abstinence normalized baseline dopamine uptake

We previously demonstrated that IntA to cocaine increases DAT efficiency during abstinence in the NAc core (Alonso et al., 2022; Clark et al., 2024). Here, we examined whether sleep restoration during abstinence mitigates these effects. Dopamine peak height and dopamine uptake were assessed in brain slices containing the NAc core using FSCV 18 h after the AD7 seeking test. Cocaine-naive rats served as controls for these experiments. Neither IntA to cocaine nor sleep restoration had a significant effect on baseline dopamine peak height relative to cocaine-naive rats. However, IntA to cocaine increased dopamine uptake and sleep restoration prevented this effect. One-way ANOVAs revealed no effect of group (*F*_(2,29)_ = 2.043, *p=*0.1479) for dopamine peak height (**Fig. 5A and B**), but there was a significant effect for dopamine uptake (*F*_(2,29)_ = 4.844, *p=*0.0153; **Fig. 5A and C**). Dunnett’s tests further revealed that IntA to cocaine significantly increased dopamine uptake compared to naive controls (*p=*0.0330) and that sleep restoration prevented this effect (*p=*0.8986).

**Figure 5.**
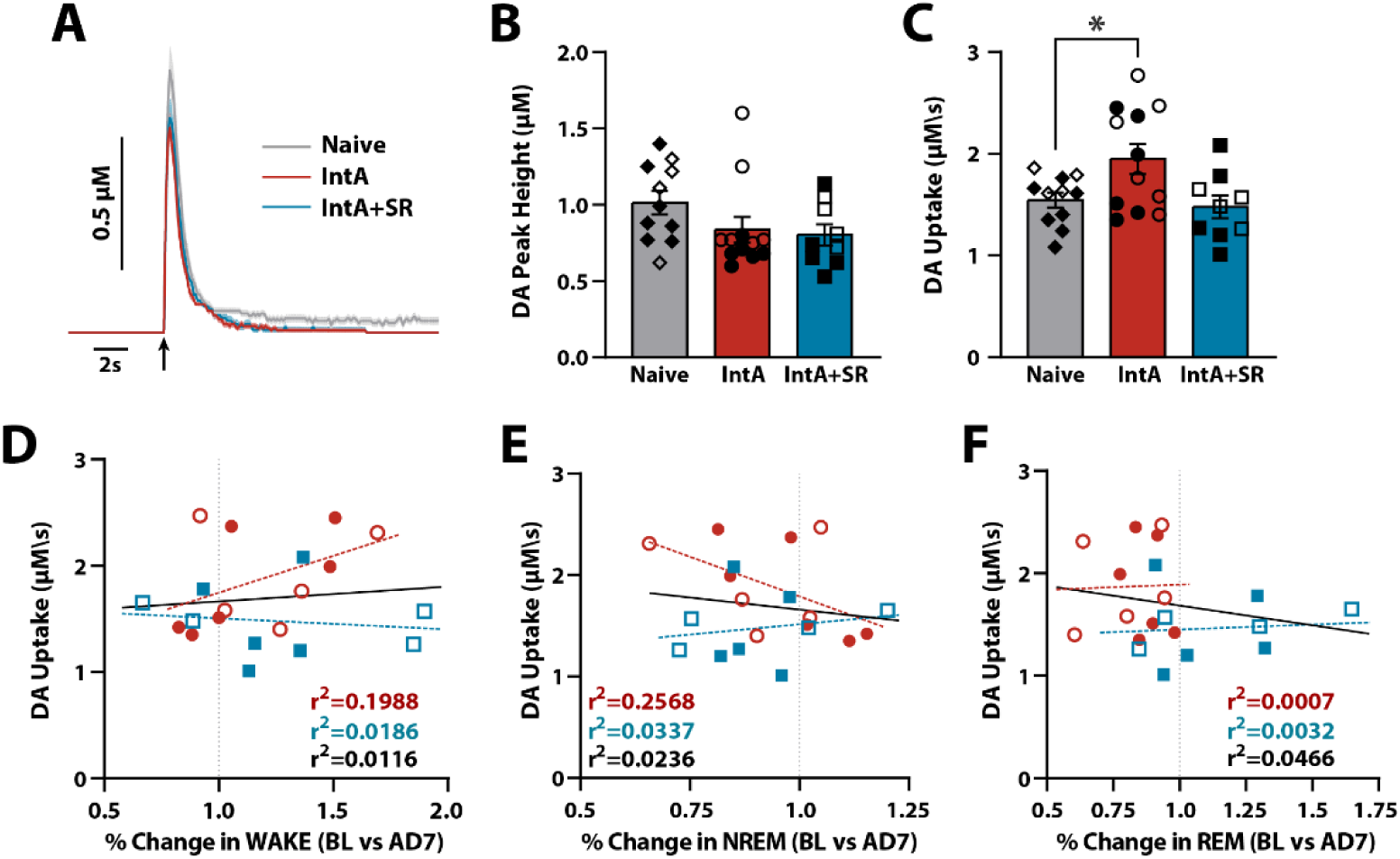
Sleep restoration during abstinence normalized dopamine uptake in the NAc core. **(A)** Average dopamine traces at baseline. Arrow indicates time of stimulation. **(B)** Dopamine peak height and **(C)** dopamine uptake at baseline (\**p*<0.05). Pearson correlations between dopamine uptake and the percentage change from baseline (BL) in time spent in **(D)** WAKE, **(E)** NREM, or **(F)** REM. Black lines depict trendline for correlations with the IntA and IntA+SR groups combined. Vertical dashed lines indicate 100% of baseline. Data shown as mean ± SEM. ◇○□females, ◆●◼ males. Naive n=11, IntA n=12, IntA+SR n=9. Dunnett’s tests \**p*<0.05 vs Naive.

We next examined if changes in sleep/wake behavior predicted the rate of dopamine uptake. A Pearson correlation for all rats (IntA and IntA+SR) indicated no significant relationship between the percentage of change in WAKE (baseline vs AD7) and dopamine uptake (*F*_(1,18)_ = 0.2122; r^2^ = 0.0116, *p*=0.6505; **Fig. 5D**). Likewise, we did not observe significant correlations for the IntA (*F*_(1,9)_ = 2.233; r^2^ = 0.1988, *p*=0.1693) or the IntA+SR groups (*F*_(1,7)_ = 01325; r^2^ = 0.0186, *p*=0.7266) when examined separately (**Fig. 5D**). When examining if the percentage of change in NREM predicted dopamine uptake we also observed no significant correlations for either all rats combined (*F*_(1,18)_ = 0.435; r^2^ = 0.0236, *p*=0.5179), for the IntA group (*F*_(1,9)_ = 3.111; r^2^ = 0.2568, *p*=0.1116), or the IntA+SR group (*F*_(1,7)_ = 0.2443; r^2^ = 0.0337, *p*=0.6383; **Fig. 5E**). Similarly, we observed no correlations between the percentage of change in REM sleep and dopamine uptake for all rats combined (*F*_(1,18)_ = 0.8810; r^2^ = 0.0466, *p*=0.3603), for the IntA alone group (*F*_(1,9)_ = 0.0065; r^2^ = 0.0007, *p*=0.9376), or the IntA+SR group (*F*_(1,7)_ = 0.0195; r^2^ = 0.0032, *p=*0.8934; **Fig. 5F**).

### Sleep restoration during abstinence normalized cocaine-induced inhibition of dopamine uptake

To assess changes in the effects of cocaine on dopamine transmission, increasing concentrations of cocaine were bath-applied to NAc core slices and changes in dopamine peak height and DAT sensitivity to cocaine (i.e. inhibition of dopamine uptake) were measured. We observed that IntA to cocaine reduced the effects of cocaine on dopamine peak height, and that sleep restoration had no effect on this measure. A mixed design ANOVA with cocaine concentration as the within-subjects variable (0.3-30 µM) and group (IntA vs IntA+SR) as the between-subjects variable revealed a significant effect of cocaine concentration (*F*_(2.308,62.31)_ = 137.9, *p*<0.0001), but no effect of group (*F*_(2,27)_ = 2.479, *p=*0.1027), or group × cocaine concentration interaction (*F*_(4.615,62.31)_ = 2.306, *p=*0.0596) for dopamine peak height (**Fig. 6A and B**). Dunnett’s tests comparing the IntA and IntA+SR groups to naive controls showed that sleep restoration decreased the effects of cocaine on dopamine peak height only at the lowest cocaine concentration (0.3μM; *p*=0.0385).

**Figure 6.**
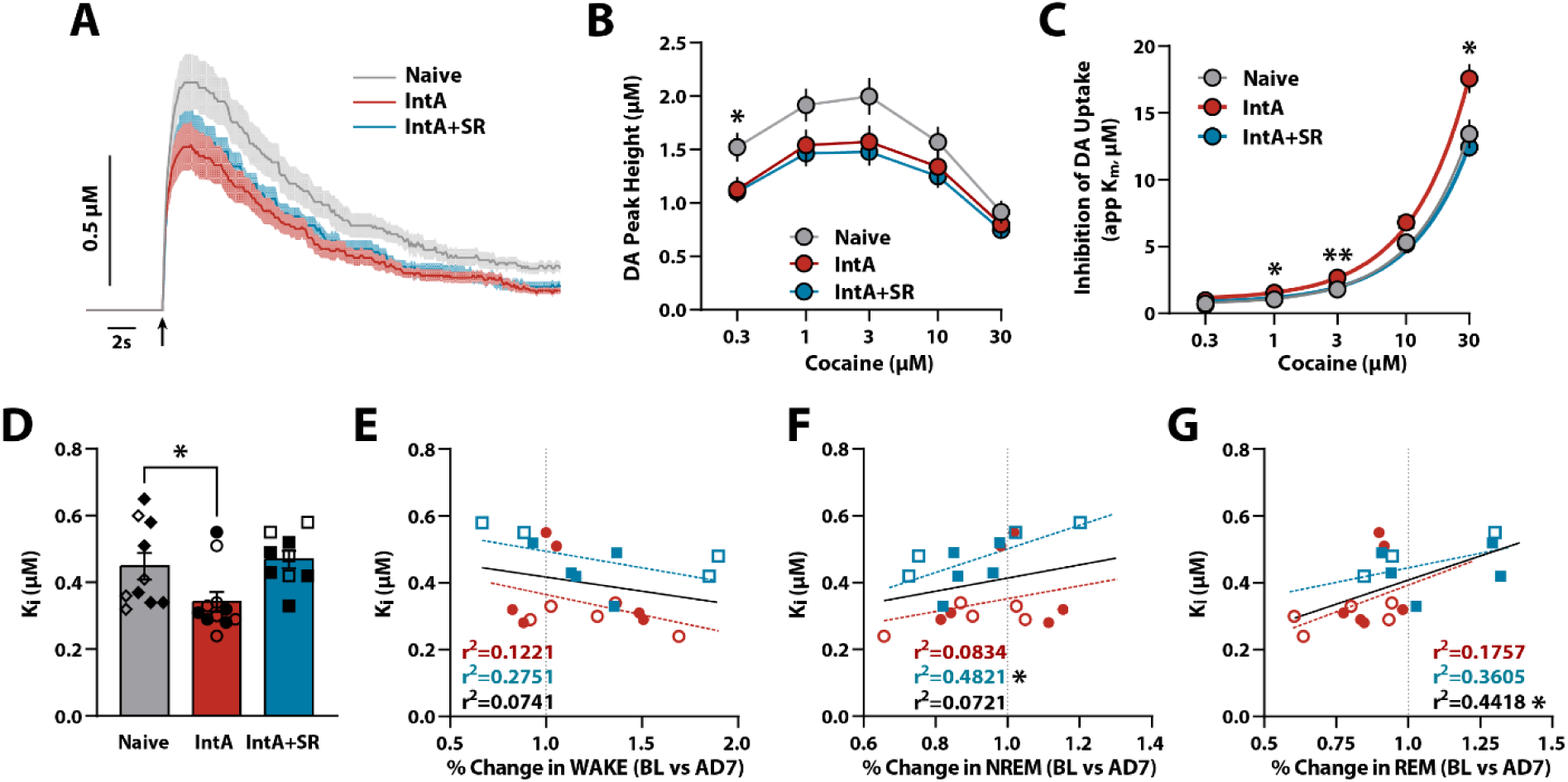
Sleep restoration normalized dopamine transporter efficiency and sensitivity to cocaine in the NAc core. **(A)** Average dopamine traces at 30μM cocaine. Arrow indicates time of stimulation. **(B)** Dopamine peak height **(C)** and inhibition of dopamine uptake across cocaine concentrations. **(D)** 50% inhibition of dopamine uptake (cocaine Ki). Pearson correlations between cocaine Ki and the percent change from baseline (BL) in time spent in **(E)** WAKE, **(F)** NREM, or **(G)** REM. Vertical dashed lines indicate 100% of baseline. Data shown as mean ± SEM. ◇○□females, ◆●◼ males. Naive n=10, IntA n=11, IntA+SR n=9. Dunnett’s tests \**p*<0.05 vs Naive. Pearson correlation \**p*<0.05.

Next, we found that IntA to cocaine increased DAT sensitivity to cocaine and sleep restoration normalized these effects. A mixed design ANOVA with cocaine concentration as the within-subjects variable (0.3-30 µM) and group (IntA vs IntA+SR) as the between-subjects variable indicated a significant effect of group (*F*_(2,27)_ = 8.620, *p=*0.0013) and concentration (*F*_(1.366,36.88)_ = 506.3, *p*<0.0001), and a group × concentration interaction (*F*_(2.732,36.88)_ = 5.864, *p=*0.0029) on DAT sensitivity to cocaine (**Fig. 6A and C**). Dunnett’s tests further revealed that IntA to cocaine significantly increased DAT sensitivity to cocaine at 1μM (*p=*0.0122), 3μM (*p=*0.0012), and 30 μM cocaine (*p=*0.0255) relative to cocaine-naive controls. However, sleep restoration prevented these changes in DAT sensitivity to cocaine. We next examined if sleep restoration influenced the cocaine concentration that produces 50% maximal uptake inhibition (Ki). A one-way ANOVA showed that a significant effect of group (*F*_(2,27)_ = 4.586, *p=*0.0193) for cocaine Ki and a Dunnett’s test further demonstrated that IntA to cocaine decreased Ki (*p*=0.0456), while sleep restoration did not (*p*=0.8675; **Fig. 6D**). Together, these observations indicate that sleep restoration attenuated the increased DAT sensitivity to cocaine observed after abstinence from IntA to cocaine.

Lastly, we examined if changes in sleep/wake behavior after IntA to cocaine predicted alterations in the effects of cocaine on dopamine transmission. A Pearson correlation for all rats (IntA and IntA+SR) indicated no significant relationship between the percentage of change in WAKE (baseline vs AD7) and cocaine Ki (*F*_(1,18)_ = 1.404; R^2^ = 0.0741, *p*=0.2457). Likewise, we did not observe significant correlations between the percentage of change in WAKE and cocaine Ki for the IntA (*F*_(1,9)_ = 1.252; R^2^ = 0.1221, *p*=0.2922), or the IntA+SR groups (*F*_(1,7)_ = 2.657; R^2^ = 0.2751, *p*=0.1471) when examined separately (**Fig. 6E**). When examining if the percentage of change in NREM predicted cocaine Ki we also observed no significant correlations for all rats combined (*F*_(1,18)_ = 1.398; R^2^ = 0.0721, *p*=0.2525), or for the IntA alone group (*F*_(1,9)_ = 0.8190; R^2^ = 0.0834, *p*=0.3891). However, there was a significant correlation between the percentage of change in NREM sleep and cocaine Ki for the IntA+SR group (*F*_(1,7)_ = 6.515; R^2^ = 0.4821, *p*=0.0380; **Fig. 6F**). Finally, we observed a significant correlation between the percentage of change in REM sleep and cocaine Ki for all rats combined (*F*_(1,18)_ = 14.25; R^2^ = 0.4418, *p*=0.0014), but this was not the case for either the IntA alone (*F*_(1,9)_ = 1.919; R^2^ = 0.1757, *p*=0.1994) or the IntA+SR groups (*F*_(1,7)_ = 3.946; R^2^ = 0.3605, *p=*0.0874; **Fig. 6G**), despite a trend for significance in the IntA+SR group. These findings suggest that IntA not only increases DAT sensitivity to cocaine in the NAc but does so in a manner that concurrently reduces REM sleep. Further, these data suggest that sleep restoration effectively normalizes DAT sensitivity to cocaine to the level of a cocaine-naive individual while promoting REM sleep during the light phase.

## DISCUSSION

In these studies, we assessed whether sleep disruptions during abstinence from cocaine are associated with incubation of cocaine seeking and dopamine terminal adaptations. IntA to cocaine followed by abstinence engendered decreased REM sleep, intensification of cocaine seeking, and increased dopamine uptake and DAT sensitivity to cocaine compared to cocaine naive rats. Interestingly, sleep restoration normalized REM sleep, prevented incubation of cocaine seeking and normalized dopamine uptake and DAT sensitivity to cocaine. These observations suggest that REM sleep disruptions during abstinence from cocaine contribute to exaggerated cocaine seeking and that DAT adaptations are a potential molecular substrate through which these changes occur.

### IntA to cocaine engenders sleep disruptions during abstinence

Previous reports indicate that abstinence from cocaine leads to time-dependent changes in sleep that persist throughout abstinence (Pace-Schott et al., 2005; Morgan and Malison, 2007; Matuskey et al., 2011; Hasler et al., 2012; Chen et al., 2015). For example, human cocaine users display increased NREM and REM sleep on the first day of cocaine abstinence and as abstinence progresses, NREM sleep begins to normalize, while REM sleep decreases (Pace-Schott et al., 2005; Morgan and Malison, 2007). Consistent with these observations, we observed an increase in NREM sleep and a trend for increased REM during the dark phase following AD1. By AD7, however, NREM sleep returned to pre-cocaine exposure levels while REM sleep disruptions continued, thus mirroring the progression of sleep disruptions observed during abstinence in clinical populations.

### Sleep restoration prevented REM sleep disruptions during abstinence

In the current studies, we used a modified sleep restoration procedure (Chen et al., 2015) that encouraged wakefulness during the dark/active phase to consolidate sleep during the light/inactive phase. Using this approach, we observed normal percentages of time spent in REM sleep by the end of the abstinence period, which appeared to be driven primarily by longer REM sleep bouts. These findings are consistent with the prior observation that sleep restoration enhanced sleep during the light/inactive phase and ultimately improved REM sleep during abstinence from cocaine (Chen et al., 2015). One limitation with the sleep restoration approach used here is that the rotating bar promotes elevated locomotor activity. Consequently, any increase in locomotor activity makes it challenging to disentangle the directs effects of sleep consolidation from possible effects due to increased locomotor activity. It is important to note that voluntary exercise (e.g. wheel running) has been demonstrated previously to attenuate incubation of cue-induced cocaine seeking (Zlebnik and Carroll, 2015; Carroll, 2021; Carroll et al., 2022), and clinical evidence suggests reduced latency to REM sleep following exercise (Driver and Taylor, 2000; Abd El-Kader and Al-Jiffri, 2020) as well as reduced REM sleep disruptions following acute sleep deprivation (Zagaar et al., 2012). However, these previous studies involve aerobic exercise, which engages large muscle groups and produces significant increases in heart rate and respiration. Based on the speed of the rotating bar in our studies, it is very unlikely that rats experienced conditions reminiscent of significant aerobic exercise. Notwithstanding, even if the effects of sleep restoration rely on increased locomotor activity, this would suggest that exercise – possibly through effects on REM sleep consolidation – is a potential behavioral therapeutic approach for reducing cocaine craving. Future studies will need to disambiguate locomotor activity from direct effects on sleep by selectively activating the bar with sleep onset in general or only for REM sleep.

### Sleep restoration attenuated incubation of cocaine seeking

The extent to which sleep disruptions following IntA to cocaine contribute to increased cocaine seeking remains unclear. Nonetheless, several observations suggest a link between disrupted sleep and motivation for cocaine. For example, sleep deprivation enhanced the acquisition of cocaine self-administration, motivation to obtain the drug, and drug seeking during cocaine abstinence (Puhl et al., 2009; Puhl et al., 2013; Chen et al., 2015). Further, exacerbating REM sleep disruptions during abstinence has been shown to accelerate the development of incubation of cocaine seeking (Chen et al., 2015). Here we confirmed that normalizing REM sleep during cocaine abstinence, attenuated cue-induced cocaine seeking, suggesting that restoring REM sleep during abstinence may serve as a behavioral therapy to attenuate cocaine craving and potentially relapse.

### Sleep restoration prevented alterations in dopamine uptake

Mesolimbic dopamine neurons are activated by exposure to drug cues during abstinence from cocaine, resulting in exaggerated motivational states that predispose individuals to seek drug and leading to relapse vulnerability (Grimm et al., 2002; Lu et al., 2004; Kawa et al., 2019). Greater responsiveness to cocaine cues following abstinence from cocaine has been associated with increased phasic dopamine signals (Willuhn et al., 2012; Ostlund et al., 2014; Saddoris et al., 2016). Additionally, presentation of cocaine-associated cues during seeking tests yields cue-evoked dopamine release within the NAc (Weber et al., 2024; Burgeno et al., 2025). However, the degree to which dopamine changes following IntA to cocaine are associated with sleep disruptions has not been sufficiently explored. In the current studies, we observed increases in dopamine uptake and greater DAT sensitivity to cocaine in NAc slices, which is consistent with our prior observations after 7 days (Alonso et al., 2022; Clark et al., 2024; Samels et al., 2024) and 28 days of abstinence (Alonso et al., 2022). Sleep restoration prevented these aberrant changes in DAT function, despite no effects on dopamine peak height observed following IntA to cocaine. These observations suggest that DAT adaptations during abstinence from IntA to cocaine may contribute to enhanced behavioral responses to cocaine as seen by the increase in cocaine seeking observed after 7 days of abstinence. Further, by preventing these DAT adaptations, sleep restoration may decrease behavioral responses to cocaine-associated cues, thus reducing cocaine seeking. Recent observations have examined the relationship between cocaine seeking behavior and cue-evoked dopamine signals in the NAc and demonstrate that both contingent and non-contingent cue presentations result in cue-evoked dopamine release during (Burgeno et al., 2023; Weber et al., 2024). However, those studies were conducted using short (1-h FR1) and long access (6-h FR1) self-administration schedules and not IntA exposure. As such, future studies will need to examine to what extent cocaine seeking following IntA to cocaine is associated with changes in cue-evoked dopamine signals.

## CONCLUSION

Our studies examined to what extent sleep disruptions during abstinence contribute to incubation of cocaine seeking and dopamine adaptations. Results demonstrated that IntA to cocaine led to decreased REM sleep, robust incubation of cue-induced cocaine seeking, increased dopamine uptake at baseline, and greater DAT sensitivity to cocaine. Importantly, sleep restoration not only normalized REM sleep and prevented the incubation of cocaine seeking, but also normalized DAT function. These findings suggest that dysregulated sleep during cocaine abstinence may be a critical factor in the persistence of cocaine craving and that interventions aimed at restoring sleep, particularly REM sleep, may serve as an effective behavioral therapy for reducing cocaine craving and preventing relapse. Moreover, our findings that sleep restoration normalized aberrant DAT adaptations observed following abstinence from IntA to cocaine suggest that the therapeutic effects of sleep restoration may be mediated through normalization of aberrant mesolimbic DAT adaptations.

## Funding Resources

This work was supported by NIH grants DA056193 and DA039100 to R.A.E. and Drexel University Dean’s Fellowship for Excellence in Collaborative or Themed Research to I.P.A.

## Acknowledgements

We thank the NIDA drug supply program for donating the cocaine hydrochloride and Bethan O’Connor for supporting the self-administration, sleep, and FSCV data collection.

## Author Contributions

IPA conceptualized and designed the studies, conducted the self-administration, sleep and FSCV data collection, analyzed and interpreted the data, wrote an original draft, and edited the content. SRC conducted self-administration, sleep and FSCV data collection, analyzed and interpreted the data, wrote an original draft, and edited the content. VMM, conducted self-administration, sleep and FSCV data collection, wrote and edited the content. RAE conceptualized and designed the studies, analyzed and interpreted the data, edited the original draft, and edited the content.

## References

1. Abd El-Kader SM, Al-Jiffri OH (2020) Aerobic exercise affects sleep, psychological wellbeing and immune system parameters among subjects with chronic primary insomnia. Afr Health Sci 20:1761–1769.

2. Alonso IP, Pino JA, Kortagere S, Torres GE, España RA (2021) Dopamine transporter function fluctuates across sleep/wake state: potential impact for addiction. Neuropsychopharmacology 46:699–708.

3. Alonso IP, O’Connor BM, Bryant KG, Mandalaywala RK, España RA (2022) Incubation of cocaine craving coincides with changes in dopamine terminal neurotransmission. Addiction Neuroscience:100029.

4. Angarita GA, Emadi N, Hodges S, Morgan PT (2016) Sleep abnormalities associated with alcohol, cannabis, cocaine, and opiate use: a comprehensive review. Addict Sci Clin Pract 11:9.

5. Angarita GA, Canavan SV, Forselius E, Bessette A, Morgan PT (2014) Correlates of polysomnographic sleep changes in cocaine dependence: self-administration and clinical outcomes. Drug Alcohol Depend 143:173–180.

6. Berridge CW, España RA (2000) Synergistic sedative effects of noradrenergic alpha(1)- and beta-receptor blockade on forebrain electroencephalographic and behavioral indices. Neuroscience 99:495–505.

7. Bjorness TE, Greene RW (2021) Interaction between cocaine use and sleep behavior: A comprehensive review of cocaine’s disrupting influence on sleep behavior and sleep disruptions influence on reward seeking. Pharmacol Biochem Behav 206:173194.

8. Borbely AA (1977) Sleep in the rat during food deprivation and subsequent restitution of food. Brain Res 124:457–471.

9. Borbely AA, Achermann P (1999) Sleep homeostasis and models of sleep regulation. J Biol Rhythms 14:557–568.

10. Brodnik ZD, Black EM, España RA (2020a) Accelerated development of cocaine-associated dopamine transients and cocaine use vulnerability following traumatic stress. Neuropsychopharmacology 45:472–481.

11. Brodnik ZD, Bernstein DL, Prince CD, España RA (2015) Hypocretin receptor 1 blockade preferentially reduces high effort responding for cocaine without promoting sleep. Behav Brain Res 291:377–384.

12. Brodnik ZD, Xu W, Batra A, Lewandowski SI, Ruiz CM, Mortensen OV, Kortagere S, Mahler SV, España RA (2020b) Chemogenetic Manipulation of Dopamine Neurons Dictates Cocaine Potency at Distal Dopamine Transporters. J Neurosci 40:8767–8779.

13. Burgeno LM, Farero RD, Murray NL, Panayi MC, Steger JS, Soden ME, Evans SB, Sandberg SG, Willuhn I, Zweifel LS, Phillips PEM (2023) Cocaine Seeking And Taking Are Oppositely Regulated By Dopamine. bioRxiv:2023.2004.2009.536189.

14. Burgeno LM, Farero RD, Murray NL, Panayi MC, Steger JS, Soden ME, Evans SB, Sandberg SG, Willuhn I, Zweifel LS, Phillips PEM (2025) Cocaine seeking and consumption are oppositely regulated by mesolimbic dopamine in male rats. Nat Commun 16:9954.

15. Calipari ES, Siciliano CA, Zimmer BA, Jones SR (2015) Brief intermittent cocaine self-administration and abstinence sensitizes cocaine effects on the dopamine transporter and increases drug seeking. Neuropsychopharmacology 40:728–735.

16. Calipari ES, Ferris MJ, Zimmer BA, Roberts DC, Jones SR (2013) Temporal pattern of cocaine intake determines tolerance vs sensitization of cocaine effects at the dopamine transporter. Neuropsychopharmacology 38:2385–2392.

17. Calipari ES, Juarez B, Morel C, Walker DM, Cahill ME, Ribeiro E, Roman-Ortiz C, Ramakrishnan C, Deisseroth K, Han MH, Nestler EJ (2017) Dopaminergic dynamics underlying sex-specific cocaine reward. Nat Commun 8:13877.

18. Carroll ME (2021) Voluntary exercise as a treatment for incubated and expanded drug craving leading to relapse to addiction: Animal models. Pharmacol Biochem Behav 208:173210.

19. Carroll ME, Dougen B, Zlebnik NE, Fess L, Smethells J (2022) Reducing short- and long-term cocaine craving with voluntary exercise in male rats. Psychopharmacology (Berl) 239:3819–3831.

20. Challasivakanaka S, Zhen J, Smith ME, Reith MEA, Foster JD, Vaughan RA (2017) Dopamine Transporter Phosphorylation Site Threonine 53 is Stimulated by Amphetamines and Regulates Dopamine Transport, Efflux, and Cocaine Analog Binding. J Biol Chem.

21. Chen B, Wang Y, Liu X, Liu Z, Dong Y, Huang YH (2015) Sleep Regulates Incubation of Cocaine Craving. J Neurosci 35:13300–13310.

22. Clark PJ, Migovich VM, Das S, Xi W, Kortagere S, España RA (2024) Hypocretin Receptor 1 Blockade Early in Abstinence Prevents Incubation of Cocaine Seeking and Normalizes Dopamine Transmission. bioRxiv:2024.2011.2030.625912.

23. Cohen SR, Xu W, Aziz NF, España RA, Kortagere S (2025) Biased Signaling Agonists of Dopamine D3 Receptor Differentially Regulate the Effects of Cocaine On Dopamine Transporter Function. ACS Chem Neurosci 16:2579–2591.

24. Dahan L, Astier B, Vautrelle N, Urbain N, Kocsis B, Chouvet G (2007) Prominent burst firing of dopaminergic neurons in the ventral tegmental area during paradoxical sleep. Neuropsychopharmacology 32:1232–1241.

25. Dolsen MR, Harvey AG (2017) Life-time history of insomnia and hypersomnia symptoms as correlates of alcohol, cocaine and heroin use and relapse among adults seeking substance use treatment in the United States from 1991 to 1994. Addiction 112:1104–1111.

26. Driver HS, Taylor SR (2000) Exercise and sleep. Sleep Med Rev 4:387–402.

27. Dugovic C, Meert TF, Ashton D, Clincke GH (1992) Effects of ritanserin and chlordiazepoxide on sleep-wakefulness alterations in rats following chronic cocaine treatment. Psychopharmacology (Berl) 108:263–270.

28. Eban-Rothschild A, Rothschild G, Giardino WJ, Jones JR, de Lecea L (2016) VTA dopaminergic neurons regulate ethologically relevant sleep-wake behaviors. Nat Neurosci 19:1356–1366.

29. España RA, Baldo BA, Kelley AE, Berridge CW (2001) Wake-promoting and sleep-suppressing actions of hypocretin (orexin): basal forebrain sites of action. Neuroscience 106:699–715.

30. Ferris MJ, España RA, Locke JL, Konstantopoulos JK, Rose JH, Chen R, Jones SR (2014) Dopamine transporters govern diurnal variation in extracellular dopamine tone. Proc Natl Acad Sci U S A 111:E2751–2759.

31. Garavan H, Pankiewicz J, Bloom A, Cho JK, Sperry L, Ross TJ, Salmeron BJ, Risinger R, Kelley D, Stein EA (2000) Cue-induced cocaine craving: neuroanatomical specificity for drug users and drug stimuli. Am J Psychiatry 157:1789–1798.

32. Garcia AN, Salloum IM (2015) Polysomnographic sleep disturbances in nicotine, caffeine, alcohol, cocaine, opioid, and cannabis use: A focused review. Am J Addict 24:590–598.

33. Grimm JW, Shaham Y, Hope BT (2002) Effect of cocaine and sucrose withdrawal period on extinction behavior, cue-induced reinstatement, and protein levels of the dopamine transporter and tyrosine hydroxylase in limbic and cortical areas in rats. Behavioural pharmacology 13:379–388.

34. Grimm JW, Hope BT, Wise RA, Shaham Y (2001) Neuroadaptation. Incubation of cocaine craving after withdrawal. Nature 412:141–142.

35. Hasler BP, Dahl RE, Holm SM, Jakubcak JL, Ryan ND, Silk JS, Phillips ML, Forbes EE (2012) Weekend-weekday advances in sleep timing are associated with altered reward-related brain function in healthy adolescents. Biol Psychol 91:334–341.

36. Kawa AB, Valenta AC, Kennedy RT, Robinson TE (2019) Incentive and dopamine sensitization produced by intermittent but not long access cocaine self-administration. Eur J Neurosci 50:2663–2682.

37. Koob GF, Le Moal M (1997) Drug abuse: hedonic homeostatic dysregulation. Science 278:52–58.

38. Levy KA, Brodnik ZD, Shaw JK, Perrey DA, Zhang Y, España RA (2017) Hypocretin receptor 1 blockade produces bimodal modulation of cocaine-associated mesolimbic dopamine signaling. Psychopharmacology (Berl).

39. Lu L, Grimm JW, Hope BT, Shaham Y (2004) Incubation of cocaine craving after withdrawal: a review of preclinical data. Neuropharmacology 47 Suppl 1:214–226.

40. Matuskey D, Pittman B, Forselius E, Malison RT, Morgan PT (2011) A multistudy analysis of the effects of early cocaine abstinence on sleep. Drug Alcohol Depend 115:62–66.

41. Meunier CJ, Roberts JG, McCarty GS, Sombers LA (2017) Background Signal as an in Situ Predictor of Dopamine Oxidation Potential: Improving Interpretation of Fast-Scan Cyclic Voltammetry Data. ACS Chem Neurosci 8:411–419.

42. Morgan PT, Malison RT (2007) Cocaine and sleep: early abstinence. ScientificWorldJournal 7:223–230.

43. Neuhaus HU, Borbely AA (1978) Sleep telemetry in the rat. II. Automatic identification and recording of vigilance states. Electroencephalogr Clin Neurophysiol 44:115–119.

44. Oishi Y, Suzuki Y, Takahashi K, Yonezawa T, Kanda T, Takata Y, Cherasse Y, Lazarus M (2017) Activation of ventral tegmental area dopamine neurons produces wakefulness through dopamine D2-like receptors in mice. Brain Struct Funct 222:2907–2915.

45. Ostlund SB, LeBlanc KH, Kosheleff AR, Wassum KM, Maidment NT (2014) Phasic mesolimbic dopamine signaling encodes the facilitation of incentive motivation produced by repeated cocaine exposure. Neuropsychopharmacology 39:2441–2449.

46. Pace-Schott EF, Stickgold R, Muzur A, Wigren PE, Ward AS, Hart CL, Clarke D, Morgan A, Hobson JA (2005) Sleep quality deteriorates over a binge--abstinence cycle in chronic smoked cocaine users. Psychopharmacology (Berl) 179:873–883.

47. Parvaz MA, Moeller SJ, Goldstein RZ (2016) Incubation of Cue-Induced Craving in Adults Addicted to Cocaine Measured by Electroencephalography. JAMA Psychiatry 73:1127–1134.

48. Phillips PE, Stuber GD, Heien ML, Wightman RM, Carelli RM (2003) Subsecond dopamine release promotes cocaine seeking. Nature 422:614–618.

49. Prince CD, Rau AR, Yorgason JT, España RA (2015) Hypocretin/Orexin regulation of dopamine signaling and cocaine self-administration is mediated predominantly by hypocretin receptor 1. ACS Chem Neurosci 6:138–146.

50. Puhl MD, Fang J, Grigson PS (2009) Acute sleep deprivation increases the rate and efficiency of cocaine self-administration, but not the perceived value of cocaine reward in rats. Pharmacol Biochem Behav 94:262–270.

51. Puhl MD, Boisvert M, Guan Z, Fang J, Grigson PS (2013) A novel model of chronic sleep restriction reveals an increase in the perceived incentive reward value of cocaine in high drug-taking rats. Pharmacol Biochem Behav 109:8–15.

52. Roberts JG, Toups JV, Eyualem E, McCarty GS, Sombers LA (2013) In situ electrode calibration strategy for voltammetric measurements in vivo. Anal Chem 85:11568–11575.

53. Roncero C, Ros-Cucurull E, Daigre C, Casas M (2012) Prevalence and risk factors of psychotic symptoms in cocaine-dependent patients. Actas Esp Psiquiatr 40:187–197.

54. Saddoris MP, Wang X, Sugam JA, Carelli RM (2016) Cocaine Self-Administration Experience Induces Pathological Phasic Accumbens Dopamine Signals and Abnormal Incentive Behaviors in Drug-Abstinent Rats. J Neurosci 36:235–250.

55. Samels SB, Shaw JK, Alonso P, Black EM, España RA (2024) Hypocretin receptor 1 blockade early in abstinence reduces future demand for cocaine. bioRxiv:2024.2012.2006.627226.

56. Shaw JK, Pamela Alonso I, Lewandowski SI, Scott MO, O’Connor BM, Aggarwal S, De Biasi M, Mortensen OV, España RA (2021) Individual differences in dopamine uptake in the dorsomedial striatum prior to cocaine exposure predict motivation for cocaine in male rats. Neuropsychopharmacology.

57. Siciliano CA, Calipari ES, Ferris MJ, Jones SR (2015) Adaptations of presynaptic dopamine terminals induced by psychostimulant self-administration. ACS Chem Neurosci 6:27–36.

58. Weber SJ, Kawa AB, Beutler MM, Kuhn HM, Moutier AL, Westlake JG, Koyshman LM, Moreno CD, Wunsch AM, Wolf ME (2024) Dopamine transmission at D1 and D2 receptors in the nucleus accumbens contributes to the expression of incubation of cocaine craving. Neuropsychopharmacology 50:461–471.

59. Willuhn I, Burgeno LM, Everitt BJ, Phillips PE (2012) Hierarchical recruitment of phasic dopamine signaling in the striatum during the progression of cocaine use. Proc Natl Acad Sci U S A 109:20703–20708.

60. Wise RA, Gingras MA, Amit Z (1996) Influence of novel and habituated testing conditions on cocaine sensitization. Eur J Pharmacol 307:15–19.

61. Woolverton WL, Johnson KM (1992) Neurobiology of cocaine abuse. Trends Pharmacol Sci 13:193–200.

62. Yang SL, Han JY, Kim YB, Nam SY, Song S, Hong JT, Oh KW (2011) Increased non-rapid eye movement sleep by cocaine withdrawal: possible involvement of A2A receptors. Arch Pharm Res 34:281–287.

63. Yorgason JT, España RA, Jones SR (2011) Demon Voltammetry and Analysis software: Analysis of cocaine-induced alterations in dopamine signaling using multiple kinetic measures. J Neurosci Methods.

64. Zagaar M, Alhaider I, Dao A, Levine A, Alkarawi A, Alzubaidy M, Alkadhi K (2012) The beneficial effects of regular exercise on cognition in REM sleep deprivation: behavioral, electrophysiological and molecular evidence. Neurobiol Dis 45:1153–1162.

65. Zimmer BA, Oleson EB, Roberts DC (2012) The motivation to self-administer is increased after a history of spiking brain levels of cocaine. Neuropsychopharmacology 37:1901–1910.

66. Zlebnik NE, Carroll ME (2015) Prevention of the incubation of cocaine seeking by aerobic exercise in female rats. Psychopharmacology (Berl) 232:3507–3513.

